# A novel in-cell ELISA assay allows rapid and automated quantification of SARS-CoV-2 to analyse neutralizing antibodies and antiviral compounds

**DOI:** 10.1101/2020.06.05.135806

**Authors:** Lara Schöler, Vu Thuy Khanh Le-Trilling, Mareike Eilbrecht, Denise Mennerich, Olympia E. Anastasiou, Adalbert Krawczyk, Anke Herrmann, Ulf Dittmer, Mirko Trilling

## Abstract

The coronavirus disease 2019 (COVID-19) caused by the severe acute respiratory syndrome coronavirus 2 (SARS-CoV-2) is currently the most pressing medical and socioeconomic challenge. Constituting important correlates of protection, determination of virus-neutralizing antibodies (NAbs) is indispensable for convalescent plasma selection, vaccine candidate evaluation, and immunity certificates. In contrast to standard serology ELISAs, plaque reduction neutralization tests (PRNTs) are laborious, time-consuming, expensive, and restricted to specialized laboratories. To replace microscopic counting-based SARS-CoV-2 PRNTs by a novel assay exempt from genetically modified viruses, which are inapplicable in most diagnostics departments, we established a simple, rapid, and automated SARS-CoV-2 neutralization assay employing an in-cell ELISA (icELISA) approach.

After optimization of various parameters such as virus-specific antibodies, cell lines, virus doses, and duration of infection, SARS-CoV-2-infected cells became amenable as direct antigen source for quantitative icELISA. Using commercially available nucleocapsid protein-specific antibodies, viral infection could easily be quantified in human and highly permissive Vero E6 cells by icELISA. Antiviral agents such as human sera containing NAbs or antiviral interferons dose-dependently reduced the SARS-CoV-2-specific signal. Applying increased infectious doses, the icNT was superior to PRNT in discriminating convalescent sera with high from those with intermediate neutralizing capacities.

The SARS-CoV-2 icELISA test allows rapid (<48h in total, read-out in seconds) and automated quantification of virus infection in cell culture to evaluate the efficacy of NAbs as well as antiviral drugs, using reagents and equipment present in most routine diagnostics departments. We propose the icELISA and the icNT for COVID-19 research and diagnostics.

## Introduction

By the time of writing, more than 6.3 million people experienced a laboratory confirmed infection with the severe acute respiratory syndrome coronavirus (SARS-CoV)-2 and more than 378,000 people died while having coronavirus infectious disease 19 (COVID-19). First surveillance studies and calculations of excess mortality rates indicate that the precise number of infections and the true number of fatalities exceed above-mentioned numbers by far.

Coronaviruses (CoVs) are positive strand RNA viruses widespread among various vertebrate hosts including bats and rodents (*1*). Together with four seasonal human CoVs (hCoVs) as well as the two other emerging hCoVs SARS-CoV-1 and MERS-CoV, SARS-CoV-2 is the seventh hCoV causing widespread human diseases (*2*). In December 2019, SARS-CoV-2 was first recognized in the Hubei province in China (*3*) from where it rapidly spread throughout the world. In addition to its genetic similarity, SARS-CoV-2 shares some clinical characteristics with SARS-CoV-1 (*4*), but also exhibits some highly relevant particularities such as an increased spreading efficacy and the length of the course of disease (*5*). On January 31, the WHO declared the SARS-CoV-2 outbreak a Public Health Emergency of International Concern. On March 11, 2020, WHO started to denote the outbreak a global pandemic. Since its beginning, the centre of the pandemic shifted from China, via Europe and Northern Americas to Central and Southern Americas. This dynamic nature of the pandemic poses an inherent danger of repetitive local and temporal reintroduction circles. Thus, even countries which coped relatively well with the first wave must prepare in terms of diagnostics capacities for potential future re-emergences.

Most SARS-CoV-2 infections lead to mild or moderate illnesses. However, a considerable fraction of cases proceeds to severe pneumonia or life-threatening acute respiratory distress syndrome. Elderly individuals and people with pre-existing comorbidities such as impaired immunity, chronic respiratory diseases, cardiovascular diseases, and cancer are more prone to suffer from severe COVID-19. The case fatality rate (CFR) is difficult to calculate in the midst of the pandemic. Depending on the age, CFR estimates of up to 18.4% for individuals older than 80 years and 1.38% (range 1.23 - 1.53) for the general population have been reported (*6*).

Given the extent, pace, and severity of the COVID-19 pandemic, diagnostics departments even in countries with highly developed medical systems struggle to provide sufficient and timely test capacities. Since nucleic acid-based pathogen detection, usually by quantitative RT-PCR based on naso-or oropharyngeal swabs, has a very short window of opportunity, assays detecting long-lasting immune responses such as antibodies are required to monitor virus spread and to estimate potential herd immunity.

With some delay, most infected individuals raise a detectable humoral immune response including specific immunoglobulins (Ig) M, IgA, and IgG (*7–9*). Neutralizing antibodies (NAbs) bind and abrogate the function of viral proteins such as the SARS-CoV-2 Spike (S) protein that are essential for virus entry into host cells, e.g., through recognition of the cognate receptor ACE2 (*10*). Accordingly, monoclonal NAbs exhibit strong therapeutic and prophylactic efficacies in SARS-CoV-2-infected human ACE2-transgenic mice (*11*). A recent vaccination study conducted in non-human primates identified NAbs as correlate of protection (*12*), indicating that a human SARS-CoV-2 vaccine should also be capable to elicit potent NAb responses. Additionally, monoclonal NAbs and NAb-containing hyper-immunoglobulin preparations may be applicable as treatment against COVID-19 (*13*). NAbs are also the backbone of convalescent plasma (CP) therapies (*14–16*) that are one of 7 clinical recommendations of the IDSA (*17*). Based on the havoc COVID-19 causes to the global economy, immunity passports and vaccination certificates, documenting protection through NAbs, have been discussed (e.g., (*18*)). Taken together, SARS-CoV-2-specific NAbs and their quantification appear to be of central importance for the medical and socio-economic management of the pandemic.

Different types of neutralization tests (NT) have been developed for SARS-CoV-2. However, to our knowledge, these assays rely on laborious microscopic counting of virus plaques or antibody-stained foci by trained personnel (*19–21*) or on genetically modified viruses such as transgenic SARS-CoV-2 mutants (*22*) or pseudo-typed viruses (e.g., Vesicular stomatitis virus [VSV] or Human immunodeficiency virus [HIV]) expressing the S protein of SARS-CoV-2 (*23, 24*). Genetically modified viruses are generally prohibited in routine diagnostics laboratories and usually not applicable in less developed regions. Due to the central importance of NAbs and the limitations of the currently available methodology, we established a cheap, simple, fast, reliable, and automatable in-cell ELISA (icELISA)-based icNT applicable in most routine diagnostics departments.

## Methods

### Cells, viruses, interferons, and sera

Caco-2 (ATCC HTB-37) and Vero E6 (ATCC CRL-1586) were cultivated in Roswell Park Memorial Institute (RPMI) and Dulbecco’s minimal essential medium (DMEM), respectively, supplemented with 10% (v/v) FCS, penicillin, and streptomycin at 37°C in an atmosphere of 5% CO. SARS-CoV-2 was isolated from a patient sample using Vero E6 and confirmed by SARS-CoV-2 diagnostic qRT-PCR. Viral titers were determined by TCID50 titration. Human IFNα2 and IFNβ were purchased from PBL Assay Science (#11101) and Peprotech (#300-02BC), respectively. The collection of serum samples has been approved by the ethics committee of the medical faculty of the University of Duisburg-Essen (20-9208-BO). Anti-SARS-CoV-2 IgG antibodies were detected using ELISA detecting SARS-CoV-2 Spike protein (Euroimmun Medizinische Labordiagnostika, Germany) according to manufacturer’s instructions

### SARS-CoV-2 icELISA and icNT

Defined doses of SARS-CoV-2 were incubated with different serum dilutions for 1 h at 37°C prior to Vero E6 infection. Neutralizing capacities were evaluated by icELISA. A detailed icELISA/ icNT protocol is provided in Suppl. File 1. The α-N mAb1 (ABIN6952435), α-N mAb2 (ABIN6952433), α-S Ab (ABIN1031551), and POD-coupled secondary antibodies (Dianova) were used.

## Results

### Quantification of SARS-CoV-2 replication and its inhibition by antiviral compounds using icELISA

We hypothesized that virus-encoded proteins expressed by infected cells should be amenable as source of viral antigens for the detection and quantification by ELISA. We optimized the experimental conditions such as cell type, virus dose, infection period, cell fixation method, blocking reagent, as well as type and concentrations of primary and secondary antibodies (Fig. 1 and data not shown). As described in the Methods section and provided as detailed laboratory protocol in the supplementary information (Suppl. File 1), we compared different commercially available antibodies for the icELISA-based quantification of SARS-CoV-2 proteins. Based on existing data on virus entry (*10*), we infected human Caco-2 cells with graded virus doses (ranging from 0.03125 to 2 PFU/cell). At 3 days post infection (d p.i.), we fixed and stained the cells with different antibodies either recognizing the S or the N protein of SARS-CoV-2. In accordance with high expression level of the N protein (*25, 26*), certain N-specific mouse IgGs exhibited a signal-to-noise ratio favourable for icELISA (Fig. 1A). A comparison of Vero E6 and human Caco-2 cells revealed that the icELISA is applicable to different cells (Fig. 1B).

**Figure 1:**
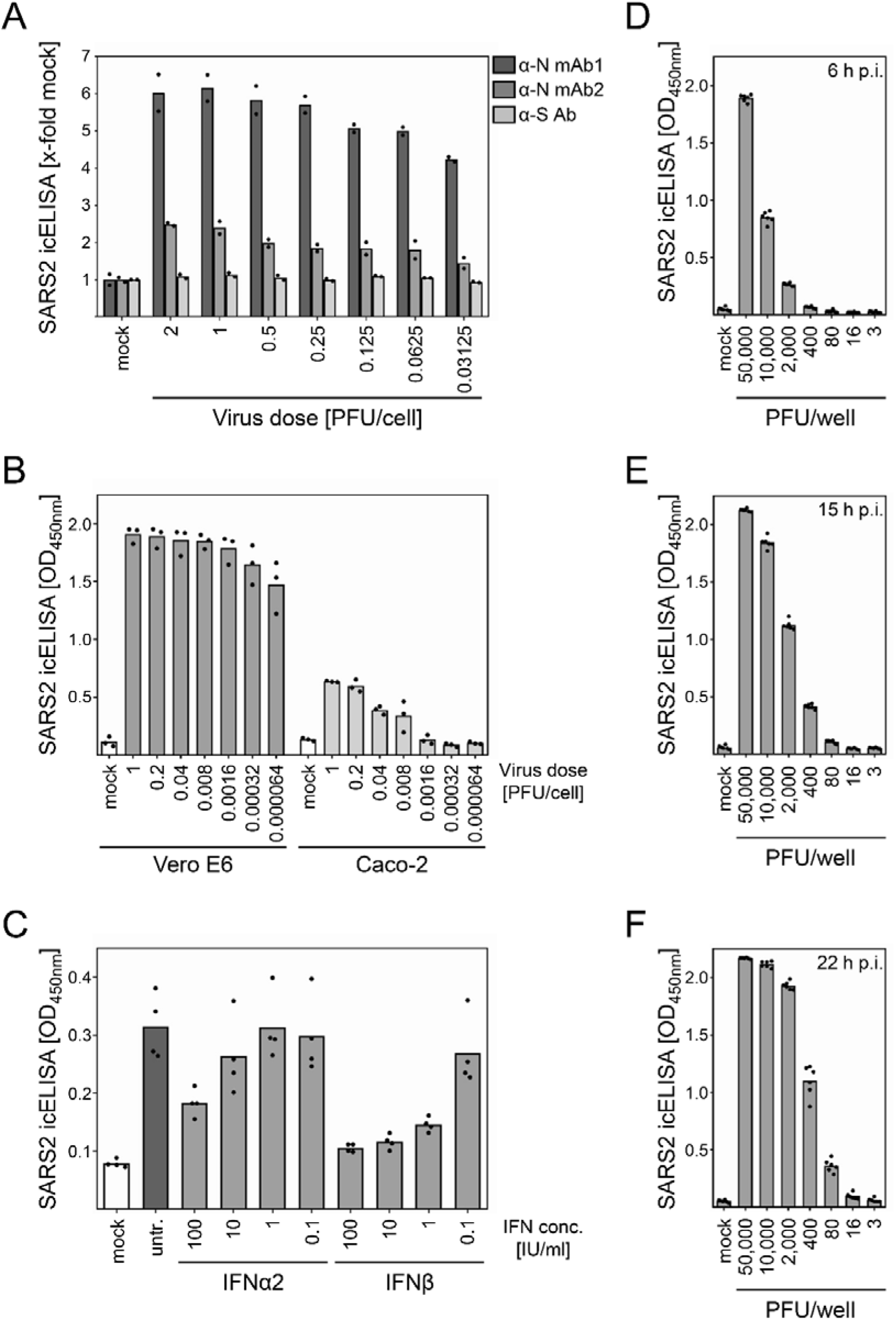
The icELISA test allows quantification of SARS-CoV-2 replication and its inhibition by antiviral compounds. (A) Caco-2 cells were infected with indicated doses of SARS-CoV-2. At 3 d p.i., cells were fixed and detected by icELISA using S-and N-specific primary antibodies. For all further icELISAs, α-N mAb1 was used. (B) Vero E6 and Caco-2 cells were infected with indicated doses of SARS-CoV-2. At 3 d p.i., cells were analysed by icELISA. (C) Caco-2 cells were treated with indicated concentrations of IFNα2 or IFNβ. At 3 h post treatment, cells were infected with SARS-CoV-2 (MOI 0.1). Viral replication was evaluated at 3 d p.i. by icELISA. (D - F) Vero E6 cells were infected with indicated doses of SARS-CoV-2. At 6, 15, and 22 h p.i. (D, E, and F, respectively), cells were analysed by icELISA. Bars depict the mean values. Dots show the values of the individual measurements.

Since the icELISA signal directly correlated with viral replication and viral antigen expression, we tested its ability to determine antiviral effects. Human Caco-2 cells were treated with graded concentrations of human interferon (IFN) α2 or IFNβ and infected 3 h later with SARS-CoV-2. At 3 d p.i., viral antigen amounts were quantified by icELISA. In accordance with a recent clinical phase 2 trial (*27*), IFNβ exhibited strong and dose-dependent antiviral activity against SARS-CoV-2 in human cells (Fig. 1C), indicating that the icELISA is applicable for future experiments addressing the efficacy of potential antiviral drugs in human cells.

Despite different start MOIs, similar icELISA signals were observed at 72 h p.i. in Vero E6 (Fig. 1B), indicating multiple rounds of virus amplification and extraordinary fast replication kinetics of SARS-CoV-2 in Vero E6 cells, consistent with previous studies (*28*). To test if shorter infection periods might result in virus dose-dependent signals, we analysed infected Vero E6 cells after 6, 15, and 22 h. SARS-CoV-2 was readily detectable in Vero E6 cells by icELISA already at 6 h p.i. (Fig. 1D-F).

Taken together, these data indicate that the icELISA allowed rapid identification and quantification of SARS-CoV-2 replication in Vero E6 and human cells as well as its inhibition by antiviral compounds.

### Neutralization tests based on icELISA allow the quantification of SARS-CoV-2 NAbs as early as 6 hours post infection

Since the icELISA allowed simple and automated quantification of SARS-CoV-2-dependent antigen expression, we tested if an icELISA-based neutralization test (icNT) can be established. We infected Vero E6 cells for 6, 15, and 24 h with graded SARS-CoV-2 infection doses (500, 5000 or 50,000 PFU/well) in the absence or presence of 2 convalescent sera in 3 different concentrations (1/20, 1/40, and 1/80 dilution). Using the high infectious dose, viral antigens became dose-dependently detectable by icELISA as early as 6 h p.i. (Fig. 2A). Both immune sera strongly and dose-dependently neutralized the virus-induced signal (Fig. 2B). At 15 h p.i., the intermediate infectious virus dose also became detectable (Fig. 2C) and both human sera efficiently neutralized the infection (Fig. 2D). At 24 h p.i., all infection conditions resulted in a strong icELISA signal (Fig. 2E), indicating that infectious doses as low as 500 PFU/well are detectable by icELISA after 24 h of infection. Even the strong icELISA signal at 24 h p.i. was dose-dependently neutralized by both sera (Fig. 2F). Please note that the neutralizing capacity of given sera dilutions was less pronounced at higher virus doses as compared to lower virus doses, as indicated by residual icELISA signals.

**Figure 2:**
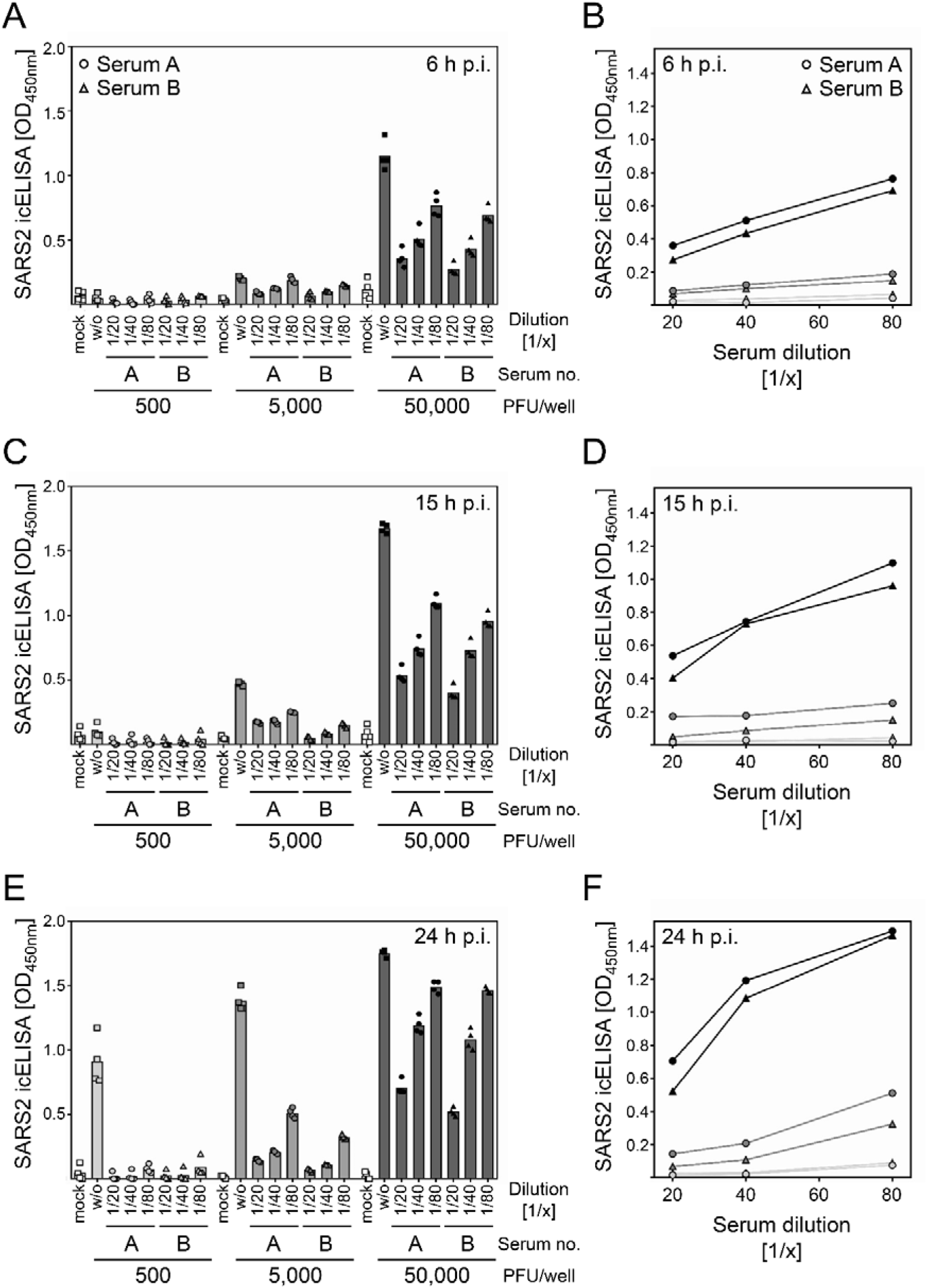
The icNT allows the quantification of SARS-CoV-2 NAbs as early as 6 h p.i. (A - F) 500, 5000, and 50,000 PFU of SARS-CoV-2 were incubated with indicated dilutions of 2 convalescent sera for 1 h before Vero E6 cells were infected. At 6 h p.i. (A, B), 15 h p.i. (C, D), and 24 h p.i. (E, F), cells were analysed by icELISA to evaluate the neutralizing capacity of the sera. (A, C, E) Bars depict the mean values. Dots show the values of the individual measurements. (B, D, F) All mean values of the different serum dilutions and virus doses were depicted in one diagram to compare the influence of the input virus amount on the course of the curve. Light grey, 500 PFU. Grey, 5000 PFU. Dark grey, 50,000 PFU.

Taken together, the icELISA resulted in a time- and virus dose-dependent signal constituting a surrogate for SARS-CoV-2 infection and replication. The fact that the infection and the resulting icELISA signal were neutralized by NAbs present in immune sera indicated that the fast and automated icELISA format is applicable for icNTs.

### The icNT results correlate with the standard SARS-CoV-2 neutralization test conducted by staining of virus foci and microscopic counting

Although most SARS-CoV-2 NTs have not been formally validated and certified, classic plaque reduction neutralization tests (PRNT) are currently considered to represent the gold standard for the detection of SARS-CoV-2-specific NAbs. Various commercially available IgM, IgA, and IgG ELISAs have been compared to PRNTs (e.g., (*28*). To validate the novel icNT, 20 sera −10 positive for SARS-CoV-2-specific IgG as determined using the Euroimmun ELISA (ELISA ratio: 1.26 to 11.39) and 10 ELISA-negative sera (ELISA ratio < 0.9) -were compared side-by-side by icNT and standard PRNT using 200 PFU/well (Fig. 3A). One set was processed by icNT, while for the other set virus foci were stained using an antibody-based AEC staining method and manually counted by microscopy. The icNT correctly recognized all 10 positive and all 10 negative sera (Fig. 3B), indicating optimal sensitivity and specificity. In total, both methods gave almost congruent results in terms of the highest dilution resulting in 50% neutralization (Pearson’s correlation coefficient 0.965), highlighting excellent test performances of the icNT compared to the PRNT.

**Figure 3:**
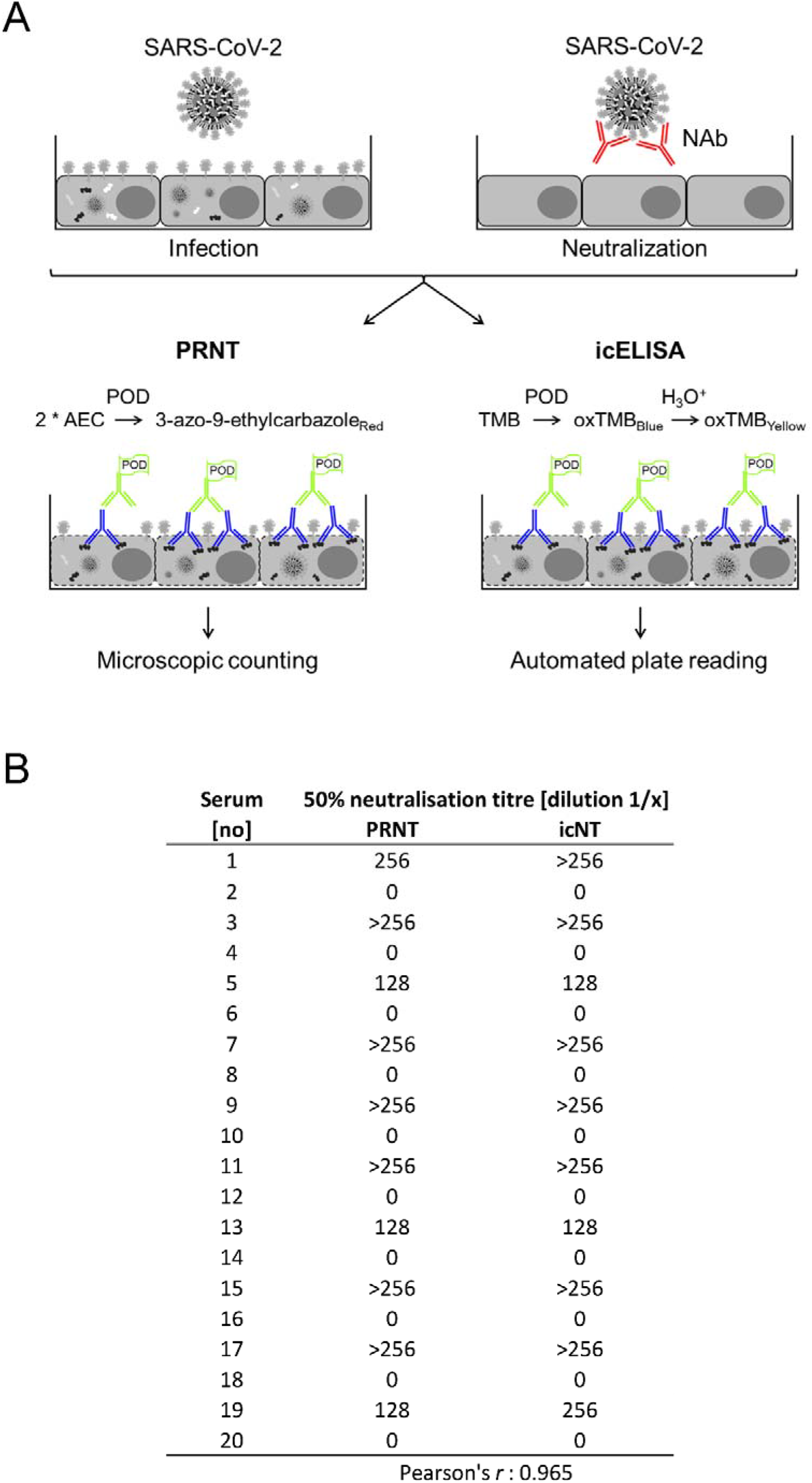
The icNT results correlate with the standard SARS-CoV-2 neutralization test. (A) Schematic representation of AEC stain-based classic NT and icELISA-based icNT. NAb, neutralizing antibody. PRNT, plaque reduction neutralization test. POD, peroxidase. AEC, 3-amino-9-ethylcarbazole. TMB, tetramethylbenzidine. (B) 200 PFU of SARS-CoV-2 were incubated with different dilutions (1/8, 1/16, 1/32, 1/64, 1/128, 1/256) of 20 serum samples (10 negatively and 10 positively tested for SARS-CoV-2 IgG by Euroimmun ELISA) for 1 h before Vero E6 were infected. At 20 and 48 h p.i., neutralizing capacity was evaluated by icELISA and AEC staining with subsequent microscopic counting, respectively. The highest dilution capable to neutralize 50% of input virus was determined and results of PRNT and icNT were compared.

### The icNT discriminates SARS-CoV-2-neutralizing sera from non-immune sera and provides superior resolution when increased virus doses are used

Standard SARS-CoV-2 NTs base on microscopic counting of plaques or antibody-stained virus foci. To enable plaque/foci recognition and individual counting by trained personnel, a countable number of PFU must be applied in PRNTs. Depending on the PRNT protocol, 100 (*19*) to 400 PFU (*20*) are used to infect each well. Based on previous experiences with virus neutralization experiments (*29, 30*), we suspected that lower infectious doses might be more susceptible to NAbs than higher virus doses, e.g., through altered ratios of NAbs and antigenic regions determining neutralization, such as the receptor binding domain (RBD) of the S protein of SARS-CoV-2. To address if the icNT can provide superior resolution in terms of discriminating intermediate from strong NAb responses in convalescent sera, we selected prototypical sera and conducted comparative icNTs applying either 200 PFU/well or 40-fold more virus (8000 PFU/well). While all positive sera clearly neutralized the low infectious dose of 200 PFU/well (Fig. 4A-C), the neutralizing capacity of the same sera considerably differed at the more restrictive high virus dose (Fig. 4D-F). Despite their neutralizing capacity at low dose infections, some of the convalescent sera (e.g., serum #19) almost did not show appreciable neutralization in high MOI infection and did not reach the typical benchmark of 50% neutralization at a 1/8 serum dilution. Thus, we concluded that the icNT principle allows higher infectious doses which appear to discriminate intermediate from superior SARS-CoV-2 neutralizing antibody titres.

**Figure 4:**
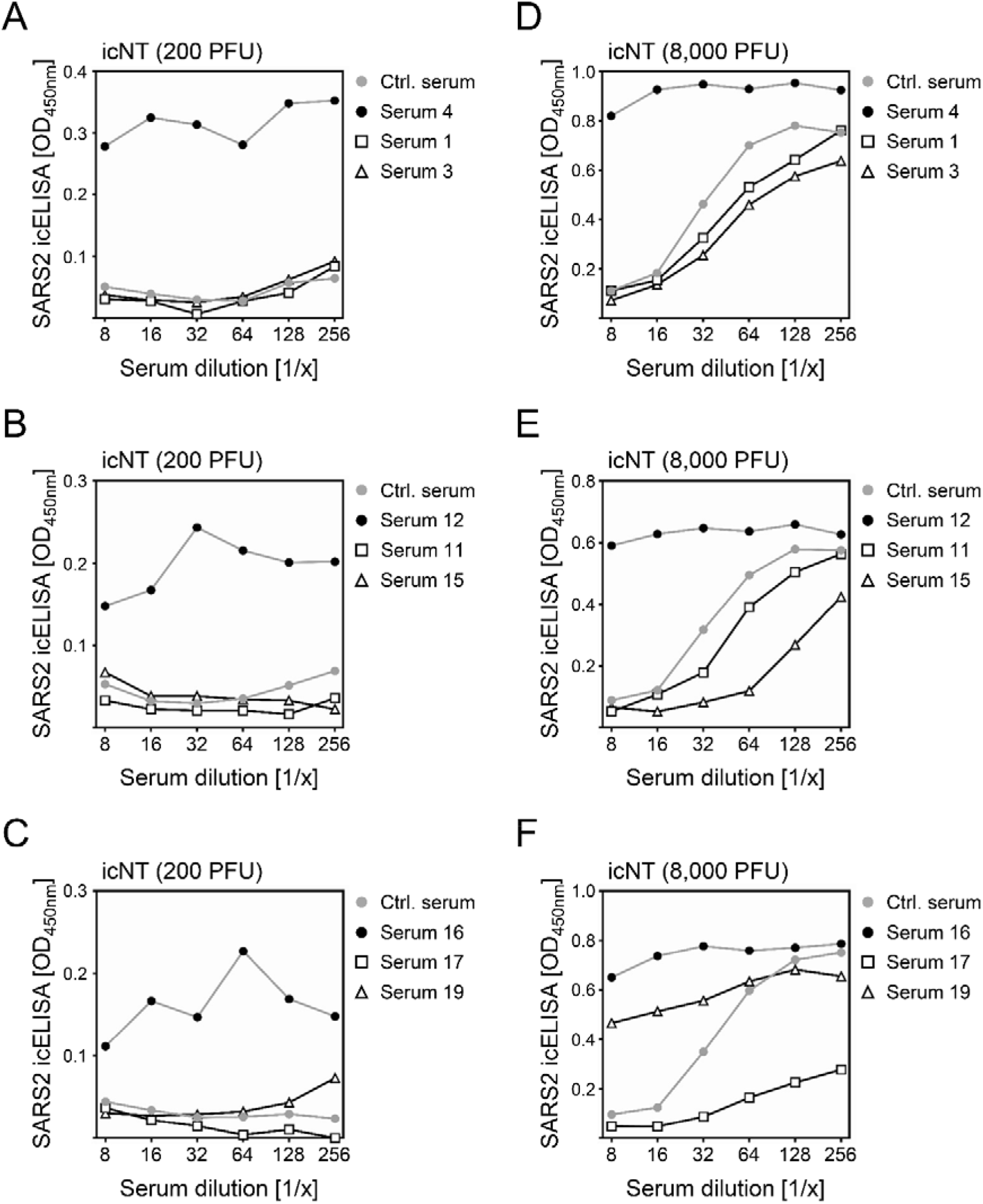
The icNT provides superior resolution upon usage of increased virus doses. (A - F) 200 or 8,000 PFU of SARS-CoV-2 were incubated with indicated dilutions of serum samples for 1 h before Vero E6 cells were infected. At 20 and 16 h p.i. (for 200 and 8,000 PFU, respectively), neutralizing capacity was evaluated by icELISA. Samples measured on the same plate are depicted in one diagram. The control serum was used as reference for all plates.

## Discussion

We established a novel icELISA-based test principle for detection and quantification of SARS-CoV-2. Given the excellent signal-to-noise ratio between infected and uninfected cells, the test was applicable to quantify the efficacy of antiviral compounds, here shown for IFNβ, and SARS-CoV-2-specific NAbs present in immune sera.

Compared to icELISA and icNT, standard virus titrations and PRNTs are far more laborious, time consuming, and expensive - not to speak from subjective and expectation biases upon usage for research. The entire icNT can be processed in less than 2 days including the infection period. Given that the protocol includes an early fixation step using 4% (w/v) paraformaldehyde which inactivates SARS-CoV-2, all subsequent steps can be processed outside the biosafety level 3 laboratory. The actual data acquisition is conducted within seconds, using standard ELISA plate readers present in most routine diagnostics departments. The icELISA and icNT provide increased data quality and precision by generating continuous data sets. Since the detection antibodies can be applied in icELISA and icNTs in relatively high dilutions (1/5000 and 1/2000 of primary and secondary antibody, respectively), the assay is relatively cheap (in our case, around 0.10 € per well for both antibodies and the TMB peroxidase ELISA substrate). The specificity of the icELISA and icNT is provided by the defined SARS-CoV-2 added to the cell cultures on purpose. The primary and secondary detection antibodies just serve to visualize and quantify viral antigens. Thus, SARS-CoV-2-specific antibodies can be applied for icELISA detection notwithstanding potential cross-recognition of other CoVs such as hCoV-HKU1 or hCoV-OC43 - simply because these viruses are not present in the culture. Obviously, such antibodies recognizing conserved residues cannot be used for classic antigen-recognizing ELISAs due to their inability to discriminate coronaviruses. More virus (>1 PFU/cell) can be applied to icNTs. Such high-PFU icNTs scrutinize the virus-neutralizing capacity of sera more strictly, enabling a higher resolution compared to PRNT assays which all rely on low virus numbers. It is tempting to speculate that sera exhibiting superior neutralization in high PFU icNT might be more beneficial in CP therapies.

NTs based on pseudo-typed viruses using heterologous expression of the SARS-CoV-2-encoded S by viruses may have certain advantages. However, genetically modified viruses are inapplicable by law in various routine diagnostics departments and usually unavailable in less developed countries. Without the use of genuine infectious viruses, assays relying on pseudo-typed viruses do not fully interrogate the full spectrum of antiviral effects for example if other viral proteins influence the system, e.g., by complement activation (*31*).

Taken together, we propose the icELISA and icNT for the quantification of SARS-CoV-2 replication and its inhibition by NAbs and antiviral compounds. By changing the detection antibody, the test principle is transferable to all other viruses.

## Acknowledgments

V.T.K.L.-T. and L.S. contributed equally to this work. We thank Benjamin Katschinski and Kerstin Wohlgemuth for excellent technical assistance. M.T. was supported by the Stiftung Universitätsmedizin Essen, Kulturstiftung Essen, the Else-Kröner Promotionskolleg ELAN, and the Deutsche Forschungsgemeinschaft (DFG) through grants RTG 1949/2, TR1208/1-1, and TR1208/2-1.

## Disclaimer

The funders had no role in study design, data collection and analysis, decision to publish, or preparation of the manuscript. The authors have no conflicts of interest.

## Author’s contributions

L.S., V.T.K.L-T., M.E., and D.M. performed research. O.E.A., A.K., A.H. provided essential reagents. L.S., V.T.K.L-T., and M.T. analysed data. L.S., V.T.K.L-T., U.D. and M.T. interpreted data. V.T.K.L-T and M.T. supervised the project. L.S., V.T.K.L-T., and M.T. wrote the manuscript. All authors reviewed the manuscript.

## SARS-CoV-2 icNT (steps 1 to 13) and icELISA (steps 6 to 13)

Materials:

- Vero E6 cells
- DMEM + Glucose/Glutamine/Pyruvate supplemented with 10% FCS, Pen/Strep
- Virus stock SARS-CoV-2 (titer: at least 2x 10^5 PFU/ml)
- Primary and secondary antibody (α-N mAb, α-mouse lgG POD-coupled)
- Paraformaldehyde (PFA)
- 1x PBS (PBS), 2x PBS
- Triton-X-100
- Tween-20
- Fetal Calf Serum (FCS)
- Tetramethylbenzidine (TMB)
- 0.5 M HCl
- Distilled or deoinized water

Equipment:

- 96-well microplate
- 37°C CO_2_ incubator
- Multichannel pipette
- Microplate reader

Notes:

- The wells of the microplate should not be allowed to dry at any point during the assay procedure.
- It is recommended to use a plate shaker during the incubation steps of the icELISA.

| REAGENT | REAGENT PREPATION INSTRUCTION | WHEN TO PREPARE? |
| --- | --- | --- |
| Wash Buffer | 500µl Tween-20 + 1 liter PBS | At start of experiment |
| 8% (w/v) Paraformaldehyde Solution | 1.) Dissolve a 2x PBS tablet in 500 ml distilled or deionized water<br>2.) Add 80 gr PFA<br>3.) Use 4 M NaOH to dissolve the PFA in PBS<br>4.) Use HCl for pH adjustment (pH 7,4)<br>5.) Fill up with distilled or deionized water (1 liter) | At start of experiment (store in aliquots at -20°C) |
| Permeabilization Buffer | For one 96-well microplate:<br>Dilute 250µl Triton-X-100 in 24.75 ml PBS.<br>Mix well by vortexing. | Prepare immediately before use |
| Blocking Solution | 3% (v/v) FCS in PBS | Prepare immediately before use |
| Incubation Buffer | 1% (v/v) FCS in PBS | Prepare immediately before use |

1. Seed Vero E6 cells into 96-well microplates the day before.
  → Perform the test at least in duplicate, better in quadruplicate.
  → Include on every plate virus control and NT with control sera as references.
  a. Cells should be 80 - 100% confluent at the day of infection.
  b. Plates can be prepared outside of the BSL3 lab.
2. Dilute the control serum and the sera to be tested 1/4, 1/8, 1/16, 1/32, 1/64, 1/128, 1/256.
  → Final serum dilutions are 1/8, 1/16, 1/32, 1/64, 1/128, 1/256, 1/512 after adding the virus solution.
  → Add further dilutions if needed.
  a. Calculate the required total volume of the different dilutions (50μ1 per well) including at least 20% excess volume.
  b. Prepare double the amount of the required volume of 1/4 serum dilution in medium (to do 2-fold serial dilutions).
  c. Prepare stepwise the other dilutions.
  d. Discard the excess amount of the last dilution.
3. Prepare the virus solution (5.5 ml per 96-well microplate).
  a. Calculate the required amount of total virus (8,000 - 10,000 PFU per well in 50μ1).
  b. Prepare the virus solution by diluting the required amount of virus stock in medium.
4. Neutralisation: Incubation of virus with serum.
  a. Add virus solution to the prepared serum dilutions (50μ1 serum dilution + 50μ1 virus solution per well).
  b. Incubate for 60 - 90 min at 37 °C.
5. Infection.
  a. Aspirate the medium of the Vero E6 cells in the 96-well microplate.
  b. Prevent cells from drying out during aspiration step.
  c. Pipette the virus-serum solution into the wells (100μ1 per well).
  d. Include the virus control and the control sera on every plate.
6. Fix cells to microplate.
  a. At 16 - 24 h p.i., add an equal volume of 8% paraformaldehyde solution to the wells containing culture media (= 4% paraformaldehyde solution)
  b. Incubate for at least 2 h (or overnight) at room temperature.
  → For experiments using non-BSL3 organisms, 15 min of fixation is sufficient.
7. Replace the lids with new ones. Wipe the plates thoroughly with 4% Dismozon. Plates can now be discharged from the BSL3 laboratory.
8. Gently aspirate the fixing solution from the microplate. Wash the microplate 3 times with 300μ1 PBS per well. Add 200μ1 PBS to the wells. The microplate (sealed with parafilm) can now be stored at 4° C for several days.
9. Permeabilize cells.
  a. Prepare permeabilization buffer (for one microplate: dilute 250μ1 Triton-X-100 in 24.75 ml PBS).
  b. Aspirate PBS and add 200μ1 of freshly prepared permeabilization buffer to each well.
  c. Incubate for 30 min.
10. Blocking.
  a. Aspirate permeabilization buffer and add 100μ1 of blocking solution (3% FCS in PBS) to each well.
  b. Incubate for 1-2 h.
11. Incubation with primary antibody.
  a. Prepare primary antibody (α-SARS-CoV-2-N, 1:5000) by diluting stock antibody in the required volume of incubation buffer (1% FCS in PBS).
  b. Aspirate blocking solution and add 50μ1 of antibody solution to each well.
  c. Incubate for 2 h at room temperature or overnight at 4°C (sealed with parafilm).
12. Incubation with secondary antibody.
  a. Prepare antibody solution by diluting stock antibody in the required volume of incubation buffer (1% FCS in PBS).
  b. Aspirate primary antibody solution. Wash the microplate 3 times with 250μ1 wash buffer per well.
  c. Aspirate the wash buffer and add 50μ1 of secondary antibody solution to each well.
  d. Incubate for 1-2 h at room temperature.
  e. Aspirate secondary antibody solution. Wash 4 times with 250μ1 wash buffer per well.
13. Signal measurement.
  a. Prepare the required amount of TMB and 0.5 M HCl.
  b. Aspirate the last wash. Make sure that you have completely removed the liquid.
  c. Add 100μ1 TMB (blue color development).
  d. Stop the reaction with 100μ1 of 0.5 M HCl (by the time the uninfected wells [mock] are starting to turn blue as well).
  e. Record data at 450 nm absorbance, 620 nm reference using a microplate reader.

